# SomaModules: a pathway enrichment approach tailored to SomaScan data

**DOI:** 10.1101/2025.07.30.667673

**Authors:** Julián Candia, Giovanna Fantoni, Francheska Delgado-Peraza, Nader Shehadeh, Toshiko Tanaka, Ruin Moaddel, Keenan A. Walker, Luigi Ferrucci

## Abstract

Motivated by the lack of adequate tools to perform pathway enrichment analysis, this work presents an approach specifically tailored to SomaScan data. Starting from annotated gene sets, we developed a greedy, top-down procedure to iteratively identify strongly intra-correlated SOMAmer modules, termed “SomaModules”, based on 11K SomaScan data. We generated two repositories based on the latest MSigDB and MitoCarta releases, containing more than 40,000 SOMAmer-based gene sets combined. These repositories can be utilized by any unstructured pathway enrichment analysis tool. We validated our results with two case examples: (i) Alzheimer’s Disease specific pathways in a 7K SomaScan case-control study, and (ii) mitochondrial pathways using 11K SomaScan data linked to physical performance outcomes. Using Gene Set Enrichment Analysis (GSEA), we found that, in both examples, SomaModules had significantly higher enrichment than the original gene set counterparts. These findings were robust and not significantly affected by the choice of enrichment metric or the Kolmogorov enrichment statistic used in the GSEA procedure. We provide users with access to all code, documentation and data needed to reproduce our current repositories, which also will enable them to leverage our framework to analyze SomaModules derived from other sources, including custom, user-generated gene sets.

## Introduction

SomaScan^1^ is a highly multiplexed, aptamer-based assay capable of simultaneously measuring thousands of human proteins broadly ranging from femto-to micro-molar concentrations. This technology relies on protein-capture SOMAmer (*Slow Offrate Modified Aptamer*) reagents,^2^ designed to optimize high affinity, slow off-rate, and high specificity to target proteins. These targets extensively cover major molecular functions including receptors, kinases, growth factors, and hormones, and span a diverse collection of secreted, intracellular, and extracellular proteins or domains. In recent years, SomaScan has increasingly been adopted as a powerful tool to discover biomarkers across a wide range of diseases and conditions, as well as to elucidate their biological underpinnings in proteomics and multi-omics studies.^3–10^ Over the past decade, the coverage of the assay has grown steadily in an approximately linear fashion, from about one thousand to eleven thousand proteins ^11^ in the most recent version of the assay. Similarly to other omics capable of measuring thousands of molecular features, an essential part of the downstream analysis is to perform a so-called pathway enrichment analysis, an approach that casts gene-level measurements in a broader biological context, thus allowing researchers to interpret their data in terms of a great variety of gene sets that may represent different functions, processes, components, or associations with disease. Since the pioneering efforts proposing Over-Representation Analysis^12^ and Gene Set Enrichment Analysis (GSEA),^13,14^ more than a hundred different algorithms and variants have been developed to implement pathway enrichment analysis. ^15,16^ Regardless of the method, however, pathway enrichment analysis relies on gene set repositories that contain the pathways or gene sets to be used as reference. Arguably the largest of them is the Molecular Signatures Database (MSigDB),^17^ which contains nearly 35,000 human gene sets compiled from KEGG,^18^ Reactome,^19^ Gene Ontology,^20^ the Pathway Interaction Database,^21^ WikiPathways,^22^ and many other publicly available resources. Most of these gene sets, however, have been derived from microarrays and other transcriptomics data. For lack of other alternatives, these gene sets are typically also used in the analysis of proteomics data, which is largely inadequate for a variety of reasons.

On the one hand, regulatory processes occurring after mRNA is made, for example mRNA rate of decay, accumulation of untranslated mRNA in stress granules or efficiency of translation, are known to play a substantial role in controlling steady-state protein abundance.^23^ For instance, in the context of the integrated stress response, ^24,25^ the presence of AU-rich elements in mRNA contributes significantly to processes that regulate gene expression, RNA decay, and rates of translation, making gene expression a weak predictor of downstream protein abundance. Spatial and temporal variations of mRNAs, as well as the local availability of resources for protein biosynthesis, are also known to strongly influence the relationship between protein levels and those of their coding transcripts.^26^ Despite the common practice to use mRNA expression as proxy for protein abundance, transcript levels are not sufficient to predict protein levels in many scenarios, as manifested by transcript and protein abundances showing weak or even negative correlations for many genes. ^27^ On the other hand, different omics technologies are known to introduce a variety of potential sources of sampling bias that may confound functional enrichment analysis. ^28–30^ Sampling biases may be introduced, for instance, by different experimental platforms having biased representations of the underlying gene ontology structure or by detecting gene products with uneven reliability. For these reasons, we argue that pathway enrichment analysis should be performed against gene sets derived from proteomics data and, more specifically, tailored to each proteomics platform. Indeed, different proteomics platforms show substantial disagreement in protein quantification; studies comparing SomaScan, Olink, mass spectrometry, and immmuno-assays showed that abundances quantified by different methods show weak or non-significant correlations for a sizable proportion of target proteins. ^31–38^ Although SomaScan has consistently showed a remarkably low variability (with a median coefficient of variation of about 5%) and sensitivity (with 98% of the measured human proteins appearing *>* 2−fold brighter in human plasma samples compared to buffer wells),^39–41^ SOMAmer reagents bind cognate proteins in their native folded conformations and may, therefore, be sensitive to proteoform complexity, which in turn may explain the observed discrepancies between SomaScan and other proteomic platforms. In addition, pre-analytical variation effects such as sample storage, repeated freeze-thaw cycles and in-vitro hemolysis may have a different impact on protein abundances measured by different assays.^42,43^

In summary, upon noting the lack of SomaScan-specific resources for pathway enrichment analysis, our work represents, to the best of our knowledge, the first endeavor aiming to fill this gap. By using data generated with the newest 11K SomaScan assay from 2542 human plasma samples, as well as publicly available data from the previous 7K assay version, we built and validated an extension of pathway collections that takes into account strongly intra-correlated SOMAmer modules, termed “SomaModules”, derived from existing gene sets. Here, we show that, in case examples chosen to explore and validate this procedure, SomaModules indeed appear to outperform standard, transcriptomic-derived gene sets to characterize pathway enrichment. We provide two public repositories based on the latest MSigDB and MitoCarta releases, containing more than 40,000 SOMAmer-based gene sets combined. This flexible framework allows users to integrate SomaModules with GSEA and other pathway enrichment analysis tools such as IPA, g:Profiler and others.

## Experimental Section

### Cohort information

Blood samples were collected from 2542 visits made by 666 participants in the Baltimore Longitudinal Study of Aging^44^ (BLSA), a study of normative human aging established in 1958 and managed by the National Institute on Aging (NIA), National Institutes of Health (NIH). The study protocol was conducted in accordance with Declaration of Helsinki principles and was reviewed and approved by the Institutional Review Board of the NIH’s Intramural Research Program. Written informed consent was obtained from all participants. Following sample pre-processing using standard protocols, EDTA plasma vials were stored at −80^◦^C. In the BLSA, visits are scheduled at regular intervals (every four years for participants younger than 60 years old; every two years for those between ages 60 and 80, and yearly for those older than 80). The number of visits per participant included in this study ranged from 1 to 15 (median = 3, mean = 3.8) and spanned the period 1993-2024, although most of them (97.8%) were from the period 2005-2024. In this study, we analyzed a total of 15 metrics obtained from short and expanded physical performance batteries, which included repeated 6-meter walks at usual and fast speeds, narrow (20-cm wide) walks, chair stands, and semi- and full-tandem standing balance tests.

### The SomaScan assay

Plasma proteomic profiles were characterized using the 11K (v5.0) SomaScan assay (SomaLogic, Inc.; Boulder, CO), which consists of 10,776 SOMAmers that target annotated human proteins. Relative protein abundances are expressed in Relative Fluorescence Units (*RFU*), fully normalized following a standard procedure to account for nuisance effects in hybridization, sample volume and plate biases, and log_10_-transformed after normalization. For further details on the assay and the data processing pipeline, see Refs.^11,40^ The SomaScan assay was run during 2024 at NIA’s SomaParadise facility (Baltimore, MD), which is certified by the SomaLogic Authorized Site program. The median sample storage time from collection to measurement was 10 years. In a separate study, we showed negligible degradation and storage effects for most SOMAmers for samples stored up to 30 years. ^41^

### Extraction of SomaModules

SOMAmers are uniquely described by their Sequence ID (“SeqId”). Most, but not all, SOMAmers are uniquely mapped to target proteins. Using Entrez gene symbols from SomaScan annotations as target identifiers, a total of 10,776 human protein SOMAmers in the 11K (v5.0) SomaScan assay are mapped to 9608 unique Entrez gene symbols, spanning 10,923 unique SeqId-Entrez gene symbol pairs. Gene sets in MSigDB are organized in collections that can be downloaded from the repository’s website. ^45^ In this work, we used the latest version available (v2024.1.Hs), released in August 2024. Because gene symbols from SomaScan annotations may not match those used to annotate MSigDB gene sets, we used gene aliases from the HUGO Gene Nomenclature Committee^46^ to maximize the pairing between annotations. As a result of this procedure, we generated a SOMAmer-based translation of MSigDB collections, in which gene sets are expressed in terms of unique SOMAmer identifiers (SeqIds).

Using the SOMAmer-based translation of MSigDB gene sets as our starting point, we developed a procedure to expand the repository by adding so-called “SomaModules”, which are subsets derived from the original MSigDB gene sets that are tightly co-expressed in SOMAmer space. In order to illustrate the procedure, let us consider the “Cellular Senescence” gene set from the Reactome collection.^47^ Whereas this pathway is comprised of 197 genes, its SOMAmer-based translation spans 138 SeqIds based on the newest 11K SomaScan assay. Using plasma proteomic profiles from 666 BLSA participants, we calculated Pearson’s correlations of all SOMAmer pairs in this pathway (Fig. 1(a)). In cases where multiple visits per participant were available, only the first (earliest) one was used at this stage to prevent spuriously biasing the correlation estimates. Once the full correlation matrix was obtained, we generated a hierarchical clustering dendrogram, which was sequentially cut at different heights to produce from *K* = 2 to *K* = 5 clusters; the total number of clusters considered was thus Σ^K=5^_K=2_ *K* = 14. For each cluster, we calculated its size, *n*, defined as the number of SOMAmers in the cluster, and its mean correlation, *r*, defined as the average correlation across all SOMAmer pairs in the cluster. Applying size and intra-cluster correlation thresholds, we kept only clusters that satisfied *n* ≥ 10 and *r* ≥ 0.5. Then, we ranked clusters in decreasing order of size (first criterion) and mean correlation (second criterion). The top cluster was chosen as the first SomaModule. After removing from the correlation matrix all SOMAmers that belonged to that first SomaModule, the process was iterated to (potentially) discover additional, non-overlapping SomaModules. Each SomaModule was identified by a collection prefix (e.g. “R” for Reactome), followed by an incremental index that indicates the pathway of origin (e.g. “R.144” for the Cellular Senescence pathway we just described) and an incremental integer suffix that indicates whether we refer to the (parent) original gene set (e.g. “R.144.0”) or a (child) SomaModule (e.g. “R.144.1” for the first SomaModule and “R.144.2” for the second one). Fig. 1(b) shows the SOMAmer network obtained from pairwise correlations; nodes represent the 138 SOMAmers in this gene set and links represent the 5280 significant pairwise correlations (p-value *<* 0.05), where green and red indicate positive and negative correlations, respectively, and where link thickness is proportional to the correlation magnitude. The two SomaModules R.144.1 (of size 45) and R.144.2 (of size 10) are indicated by different node colors. It should be noticed that the fundamental building blocks of SomaModules are SOMAmers, uniquely identified via their SeqIds; however, by mapping them to gene identifiers, some node labels are repeated. For instance, SeqId “10346-5” and “10354-57” are two SOMAmers in SomaModule R.144.1 both mapped to STAT3. In this example, we observe that SomaModule R.144.1 includes STAT3, cyclin-dependent kinases (CDK), mitogen-activated protein kinases (MAPK), ribosomal proteins (RPS), and ubiquitins (UB); in turn, SomaModule R.144.2 is comprised of various histone complex proteins. Table 1 shows the list of all SomaModule collections derived from MSigDB currently available and their size (i.e., the number of gene sets contained in each collection). The latter, naturally, depends on the size and intra-cluster correlation thresholds used in the SomaModule extraction procedure. As pointed out by Ramanan et al, ^48^ a commonly chosen minimum threshold for pathway size is 10 genes, ^49–52^ which appears suitable to prevent false positive associations due to large single-gene or single-SNP effects.^53^ The intra-cluster correlation threshold, in turn, dictates the tradeoff between false positives (favored by lower thresholds) and false negatives (favored by higher thresholds). In this work, by imposing the *r* ≥ 0.5 constraint, we aimed at generating robust SomaModules. Despite this somewhat conservative choice, we obtained a relatively large number of SomaModules. Indeed, across all MSigDB collections, we generated 14,363 (child) SomaModules out of 27,342 (parent) gene sets that passed the *n* ≥ 10 size constraint after being translated in terms of SOMAmers; in other words, we expanded the original gene sets by slightly more than 50%. As with every other aspect of pathway analysis, however, threshold criteria should be dictated by the context and characteristics of the study, and may differ from the choices outlined in this work. For reference, Supplementary Table 1 shows the size of MSigDB collections derived from different threshold parameter choices.

**Figure 1:**
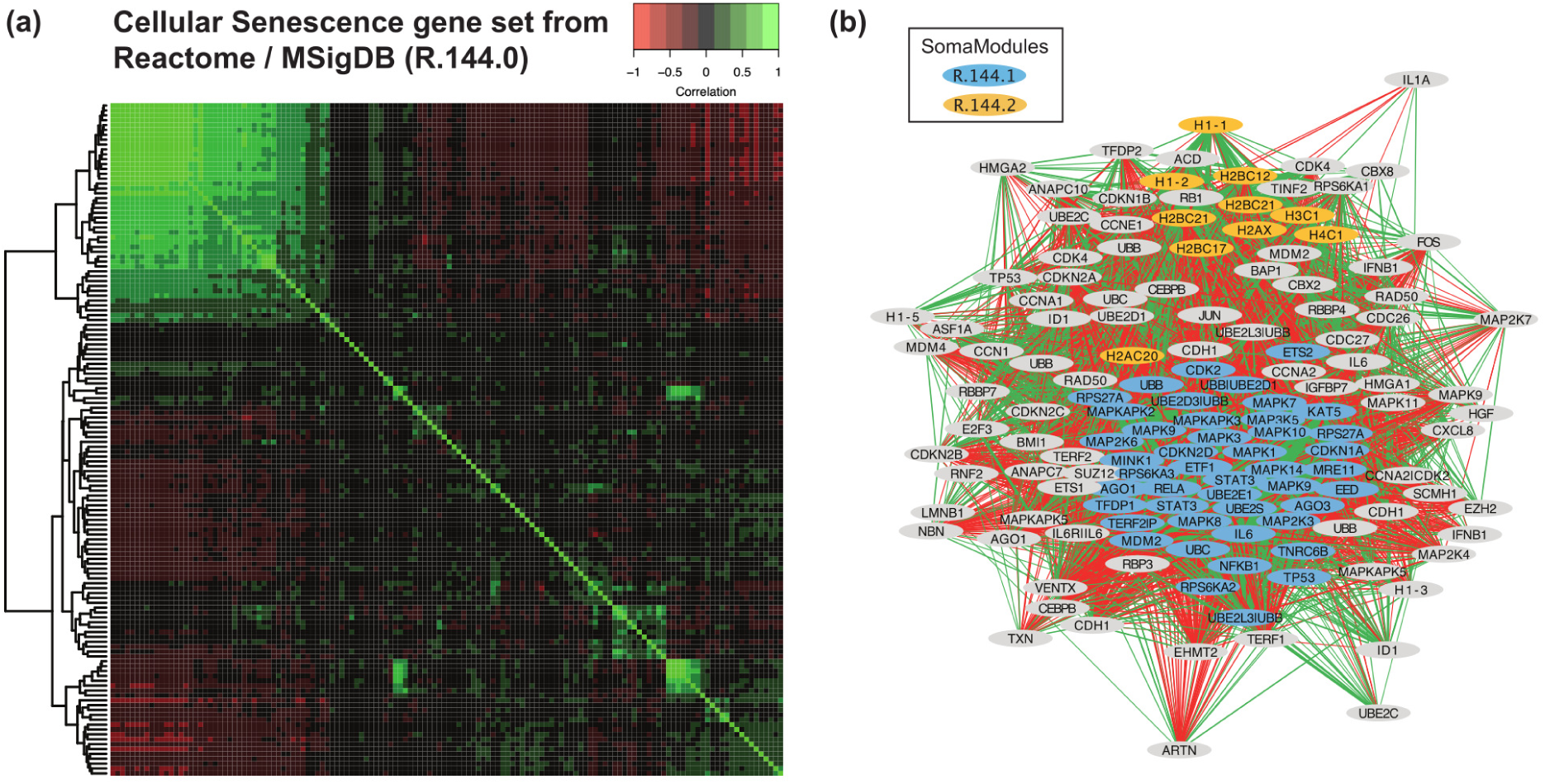
Extraction of SomaModules. **(a)** Pearson’s pairwise correlation matrix of SOMAmers mapped to the Cellular Senescence gene set from Reactome. Data were generated using the 11K SomaScan assay on plasma samples obtained from 666 BLSA participants. **(b)** SOMAmer correlation network, where green and red links represent significant positive and negative correlations, respectively. The original gene set (R.144.0) is comprised of 138 SOMAmers; our procedure identified two SomaModules, R.144.1 and R.144.2, consisting of 45 and 10 SOMAmers, respectively.

**Table 1:**
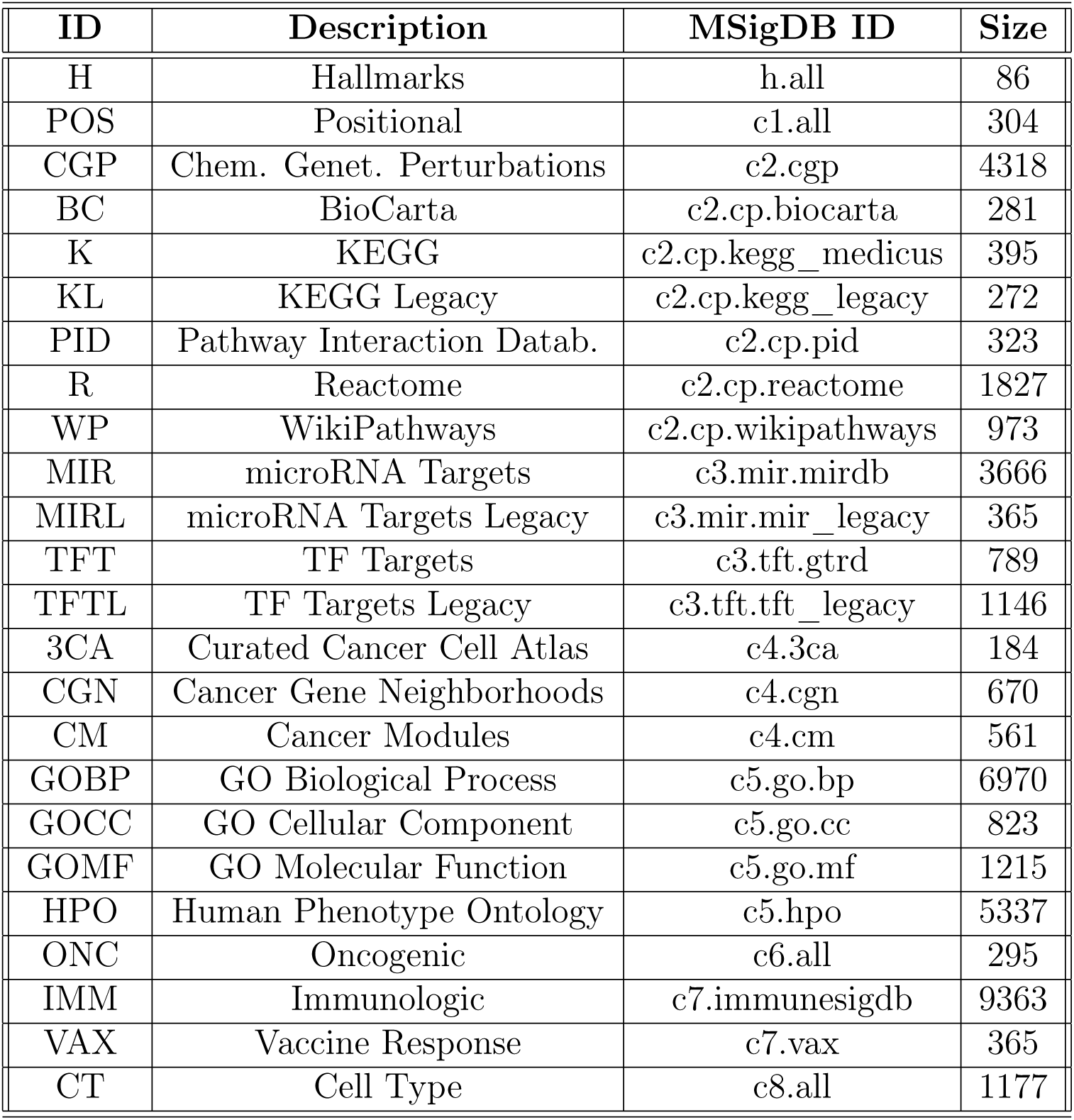
Collections from MSigDB in the SomaModule repository.

### SomaModule characteristics, limitations, and alternative approaches

The SomaModule framework is a greedy approach to identify, through successive iterations, the largest non-overlapping, strongly intra-correlated modules. In order to explore the characteristics of these modules, we calculated: (i) all SOMAmer-SOMAmer correlations within each parent gene set, (ii) those within each child SomaModule, and (iii) the cross-correlations between SOMAmers in the parent gene set (not included in the SomaModule) and those in the SomaModule. Then, for each parent-child pair, we calculated: (i) the mean correlation within the parent gene set, (ii) the mean correlation within each child SomaModule, and (iii) the mean cross-correlation between parent and child (excluding SOMAmers present in both parent and child). For the sake of simplicity, here we considered only the first child of each parent gene set, since very few gene sets have more than one child SomaModule. Supplementary Fig. 1 shows mean correlation density distributions across all parent-child pairs from different collections (Hallmarks, KEGG, Reactome and WikiPathways). As expected, all child SomaModule distributions (in red) show strong within-module correlations (*r >* 0.5 by definition). Not surprisingly, the within-parent gene set correlations (in black) are generally much weaker. Interestingly, we observe that the parent-child cross-correlation distributions (in green) are shifted towards negative values; that is, the hierarchical clustering procedure used to extract SomaModules is biased to SOMAmer clusters that are negatively correlated with the rest of the SOMAmers in the parent gene set. Indeed, this characteristic may be explained as due to the fact that SomaModules are preferentially selected to contrast with the background.

Naturally, the hierarchical-clustering-based procedure to extract SomaModules described above is just one approach out of many possible alternative formulations. For the sake of illustration, let us consider one possible use of the popular Weighted Gene Co-expression Network Analysis (WGCNA^54^) in this scenario. More specifically, let us revisit the SOMAmer correlation matrix from Fig. 1(a) and apply WGCNA to find the optimal partition of this parent gene set into submodules. WGCNA is a tool with multiple parameters to adjust. To determine the optimal power (*β*) to weigh correlations (such that increased power suppresses weaker correlations), WGCNA proposes a grid-search approach, in which a range of *β* values is explored against a scale-free topology model fit. Supplementary Fig. 2 shows that, in this case, *β* = 5 was the best fit. WGCNA offers a large variety of parameters to extract clusters. Consistent with our previous assumption, we set “minModuleSize” (the minimum module size for module detection) equal to 10. Because we wanted to discriminate up- vs down-regulated SOMAmers, we discarded the “unsigned” network type, leaving us with “signed” and “signed hybrid” modalities. Additionally, we explored the “deepSplit” parameter (which provides a simplified control over how sensitive module detection should be to module splitting) and three Topological Overlap Measure (TOM) types (“none”, “unsigned”, and “signed”). Results are summarized in Supplementary Table 2. In order to select the optimal clustering, our criterion was to maximize the mean variance explained by the first principal component (averaged over all clusters). We found that the optimal clustering consists of three clusters of sizes 88 (corresponding to the “noise” or “background” cluster), 40, and 10 SOMAmers, respectively. The latter two are fully included in the previously identified SomaModules “R.144.1” and “R.144.2”, respectively, shown in Fig. 1(b). The concordance between SomaModules and the optimal WGCNA clustering is highly significant (Fisher’s exact test p-value = 10^−42^).

Top-down clustering approaches, such as those exemplified here to find subclusters within previously annotated pathways, naturally preclude the possibility of finding de novo gene sets. This, in fact, could be achieved by implementing bottom-up procedures, similar to those used to build de-novo transcriptomic modules in blood.^55–57^ The task to extract de-novo modules presents some challenges, for instance: (i) to define criteria for the optimal number and size of clusters; (ii) to assess module stability and robustness; and (iii) to annotate denovo modules using reliable and biologically meaningful descriptors. Nonetheless, this is a promising direction of research that may complement the top-down approach we followed in this paper.

As stated in the Introduction, the goal of this work is to address potential sources of bias that may confound functional enrichment analysis, accounting for differences between transcriptomics- and proteomics-derived gene sets, as well as capturing SomaScan-specific characteristics such as proteome coverage, sensitivity, and specificity. Other sources of biological bias, caveats, and shortcomings of pathway analysis, however, remain to be carefully assessed on a case-by-case basis. One of them is referred to as biological bias, to account for the fact that cell types have highly specialized omics profiles, which can be used to reliably identify the tissue of origin. On the other hand, although disease-specific traits are typically assessed by contrast to pathway references derived from a healthy cohort, this procedure may mask disease subtypes that would require disease-specific pathway references, such as e.g. the COVID-19 Disease Map.^58^ In this regard, the Molecular Signatures Database, and therefore also the SomaModule repository derived from it, contain thousands of tissue- and disease-specific gene sets, such as those included in the Cell Type (CT), Curated Cancer Cell Atlas (3CA), Cancer Gene Neighborhoods (CGN), Cancer Modules (CM), Oncogenic (ONC) and Human Phenotype Ontology (HPO) collections (see Table 1). The choice of reference gene set collections is dictated by the context of the scientific question under study; to illustrate this point, we discuss below an application in which the relationship between physical performance and mitochondrial function in aging is investigated using SomaModules derived from a collection of annotated mitochondrial pathways.

Another challenge of pathway enrichment analysis is that of gene set overlap, where some genes participate in multiple gene sets. This phenomenon occurs in the presence of multifunctional genes (i.e., genes that play a role in several biological functions or molecular processes) but is also prevalent in some gene set collections with redundancies or a strong hierarchical structure, such as Reactome and the three lineages of Gene Ontology (GO). Beyond some early attempts to address this issue,^59–61^ this remains an important topic that deserves further investigation and lies beyond the scope of the present work.

### SomaModule repository and usage

Closely following the structure of the MSigDB public repository, we generated our first release of SomaModule collections in Gene Matrix Transposed (.gmt) file format^62^ and made them publicly available on the Open Science Framework repository. ^63^ To ensure data provenance, we followed the same naming conventions adopted by MSigDB,^45^ adding a suffix that indicates the SomaModule build version. Although we plan to maintain the SomaModule repository updated as new MSigDB versions are released, we also provide all the code, documentation and data needed to re-generate the full repository, so that users are free to explore the SomaModule framework to suit their needs and interests (for instance, by using different module selection criteria). Our framework, moreover, is not limited to MSigDB collections. To emphasize this point, we generated a second SomaModule repository from MitoCarta3.0,^64^ which contains annotated mitochondrial pathways, and used it to explore the longitudinal association between mitochondrial processes and physical performance metrics (see Results and Discussion section below). Our framework can be similarly leveraged to analyze gene sets from other sources, including custom, user-generated gene sets. Gene set collections in .gmt file format can be seamlessly uploaded into GSEA ^65^ but, simply consisting of lists of gene identifiers, they can also be used with other non-topology-based pathway analysis tools, e.g. those implementing Over Representation Analysis and Functional Class Scoring methods.^15,16^ Let us point out that GSEA does not allow gene identifiers to use hyphens (dashes), which are in fact used in SomaScan’s SeqIds; we circumvent this issue by replacing them programatically with underscores. Therefore, we provide two different versions of our repository build (using either original or modified SeqIds), as well as R scripts showing how to reformat SeqIds when uploading SOMAmer rank lists into GSEA.

### Gene Set Enrichment Analysis (GSEA)

GSEA is a well-established tool to assess pathway enrichment of ranked gene lists. ^14^ GSEA is a threshold-free method that analyzes all measured genes on the basis of their ranking score, without prior gene filtering. Let *r_k_* be the ranking score associated with the *k*-th gene, *g_k_*, with *k* = 1*, …, n* an index running through all measured genes. Without loss of generality, we assume genes to be indexed in decreasing order of their ranking scores. Typically, it is assumed that ranking scores span positive and negative values; the absolute value of a ranking score is a measure of effect size and/or statistical significance, whereas the sign indicates the direction of change (see below for further details on SOMAmer rank metrics adopted for the analysis of our case examples).

Let G*_i_* be a gene set (after removing any genes not included in the universe of measured genes). The step variable *x_k_* is defined as

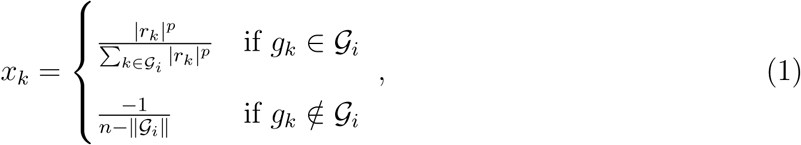

where *p* is the Kolmogorov enrichment statistic, whose values are *p* = 0 (corresponding to the classic/unweighted GSEA version) and *p* = 1, 1.5, and 2 (corresponding to weighted GSEA versions). This weight parameter was introduced to emphasize the role of genes at the top and bottom of the gene list (i.e. those with larger absolute values of the ranking score in either direction) in detriment of genes in the center of the list (i.e. those with lower absolute values of the ranking score in either direction), although recent assessments using RNA-Seq-based benchmarks suggested that the classic/unweighted approach offered comparable or better sensitivity-vs-specificity tradeoffs. ^66^ The cumulative enrichment of gene set G*_i_* is defined as the running sum of the step variable *x_k_* from top to bottom, i.e.

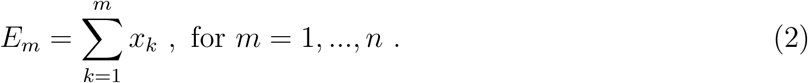

In this way, starting at *E*_0_ ≡ 0, GSEA progressively examines genes from the top to the bottom of the ranked list, increasing the cumulative enrichment if the gene at that position belongs to the pathway and decreasing the running sum otherwise. The step variable is normalized to ensure that *E_n_* = 0 at the bottom of the list. Then, the target gene set’s Enrichment Score (ES) is defined as the maximum departure from zero (in either direction, i.e. positive or negative) along the cumulative enrichment profile; its sign indicates whether the observed enrichment is associated with the top (positive ES) or bottom (negative ES) portion of the ranked gene list. In order to control for pathway size, GSEA also computes a Normalized Enrichment Score (NES). Results described in the main section of this paper used GSEA in Preranked mode with 1000 gene set permutations and the classic/unweighted Kolmogorov enrichment statistic. Supplementary Data files report results obtained with all (unweighted or weighted) choices for the Kolmogorov enrichment statistic.

### SOMAmer rank metrics

We assume a rank metric of the form

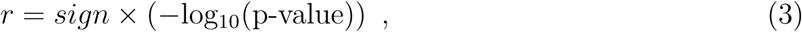

where *sign* = ±1 indicates the direction of change and p-value is an appropriate measure of the significance of that change. In the Results and Discussion section below, we explore two case examples with different types of study design. For the first one, a case-control study, we fitted SOMAmer expression values against health condition (AD or control), sex, age, and race using functions lmFit, eBayes and topTable from the R package limma v.3.56.1, which provide the AD/control fold change (log*FC*) and corresponding p-value estimated by empirical Bayes variance stabilization.^67^ Volcano plots in Supplementary Fig. 3 show the results obtained from plasma (panel (a)) and CSF (panel (b)) samples. Using *sign* (log*FC*) in Eq.(3), we obtain the rank score. For the second case example, a longitudinal analysis of physical performance, mixed-effects models were run according to the formula: “phys.perf. ∼ (1|subject)+sex+age+log_10_ (*RFU*)”, where each physical performance outcome was regressed against fixed effects for sex, age, and SOMAmer relative concentration, considering subject identifiers as random effects. The statistical significance of the SOMAmer abundance term was assessed by performing a likelihood ratio test to compare the full model against the null model: “phys.perf. ∼ (1|subject) +sex +age” using the anova function from R package stats v.4.4.0. Mixed-effects models were implemented via the lmer function from R package lme4 v.1.1.35.5 with the argument REML=FALSE to optimize the log-likelihood.^68^

### Data and Software Availability

Data analysis was performed in R v.4.4.0. Repositories in .gmt file format were imported using R package fgsea v.1.30.0. Clinical data were converted from SAS format using R package haven v.2.5.4. Fitting procedure results were parsed with R package broom v.1.0.6. GSEA runs were generated using the command-line client gsea-cli.sh v.4.3.3. Wilcoxon and t-tests were performed using base R package stats v.3.6.2. Plots were created using R packages RColorBrewer v.1.1-3, venn v.1.12, gplots v.3.1.3.1 and calibrate v.1.7.7. Anonymized datasets and custom R source code used in our analyses are available on the Open Science Framework repository at osf.io/wemcs (DOI 10.17605/OSF.IO/WEMCS).

## Results and Discussion

We will illustrate the use of SomaModules by means of two case examples: (i) a case-control proteomic analysis of Alzheimer’s Disease (AD) using the 7K SomaScan platform to examine AD-specific pathways, and (ii) a longitudinal analysis of physical performance using the 11K SomaScan assay to investigate the enrichment of mitochondrial pathways. These applications will showcase how SomaModules can be broadly used to analyze different clinical traits across different types of study design and different SomaScan versions.

Our first case example is from a cohort of control (*n* = 18) and AD (*n* = 18) patients in the Emory Goizueta Alzheimer’s Disease Research Center^69^ using the 7K SomaScan assay on plasma and cerebrospinal fluid (CSF) samples. After scanning all pathways in MSigDB, we identified six of them that were AD-specific (based on their names) and had at least one SomaModule. Fig. 2 shows Venn diagrams with the number of overlapping SOMAmers in the 7K assay based on the original MSigDB pathways (panel (a)) and the first SomaModules derived from them (panel (b)). Here, we notice that a large number of SOMAmers were specific to individual pathways; in particular, gene set WP.44 from WikiPathways and gene sets CGP.178 and CGP.181 from the Chemical and Genetic Perturbations (CGP) collection contain a large proportion of non-overlapping SOMAmers, both in their original versions (panel (a)) as well as in their SomaModule versions. Using plasma samples, Fig. 3 shows pathway enrichment quantified by means of GSEA’s Enrichment Scores (ES) and Normalized Enrichment Scores (NES), the latter of which was designed to account for differences in gene set size. Here, enrichment is defined as protein abundance in controls relative to AD patients; in agreement with an independent study, ^70^ plasma protein abundance is biased to be generally greater in controls compared with AD patients (Supplementary Fig. 3(a)). We observe that, for all six AD-specific pathways, SomaModules (red symbols) are more sensitive than the original MSigDB counterparts (blue symbols). For instance, KEGG’s Alzheimer’s Disease original gene set (KL.9.0) has NES = 1.6 and p-value ≥ 0.05, whereas the SomaModule derived from it (KL.9.1) has NES = 4.4 and p-value *<* 10^−3^. Although results shown in Fig. 3 correspond to GSEA’s classic/unweighted Kolmogorov-Smirnov enrichment statistic, we explored all weighted options (enrichment statistic parameters *p* = 1, 1.5, and 2) with similar results (Supplementary Data 1). Paired Wilcoxon and Student’s t-tests comparing the absolute value of enrichment scores between original MSigDB gene sets and SomaModules are statistically significant in all cases considered; we thus conclude that SomaModules show significantly higher enrichment than the original MSigDB gene set counterparts. In a similar vein, Fig. 4 shows pathway enrichment results obtained from CSF samples. Here, enrichment is defined in the opposite direction, as protein abundance in AD patients relative to controls (Supplementary Fig. 3(b)). Results for all GSEA weighted options are provided in Supplementary Data 2. Although only a few of the selected pathways are significant, we observe that, in cases where they are, the enrichment scores of SomaModules are greater than those of their corresponding original MSigDB counterparts.

**Figure 2:**
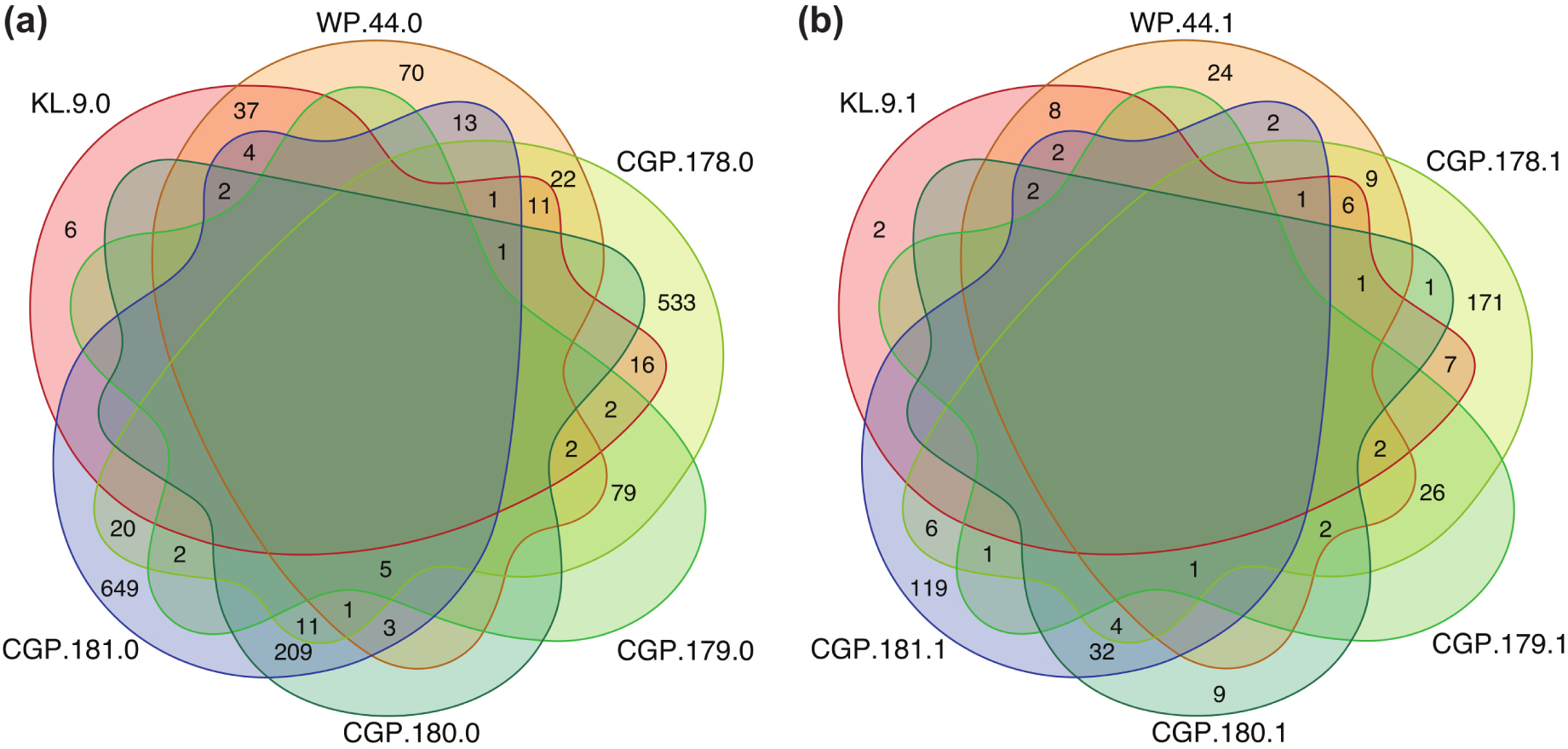
Overlaps among AD-specific pathways in the 7K SomaScan assay. Each intersection shows the number of overlapping SOMAmers; zeros are not shown. Pathways were obtained from the KEGG Legacy (KL), WikiPathways (WP), and Chemical and Genetic Perturbations (CGP) collections included in MSigDB. **(a)** Original (MSigDB) gene sets. **(b)** SomaModules derived from the original (MSigDB) gene sets.

**Figure 3:**
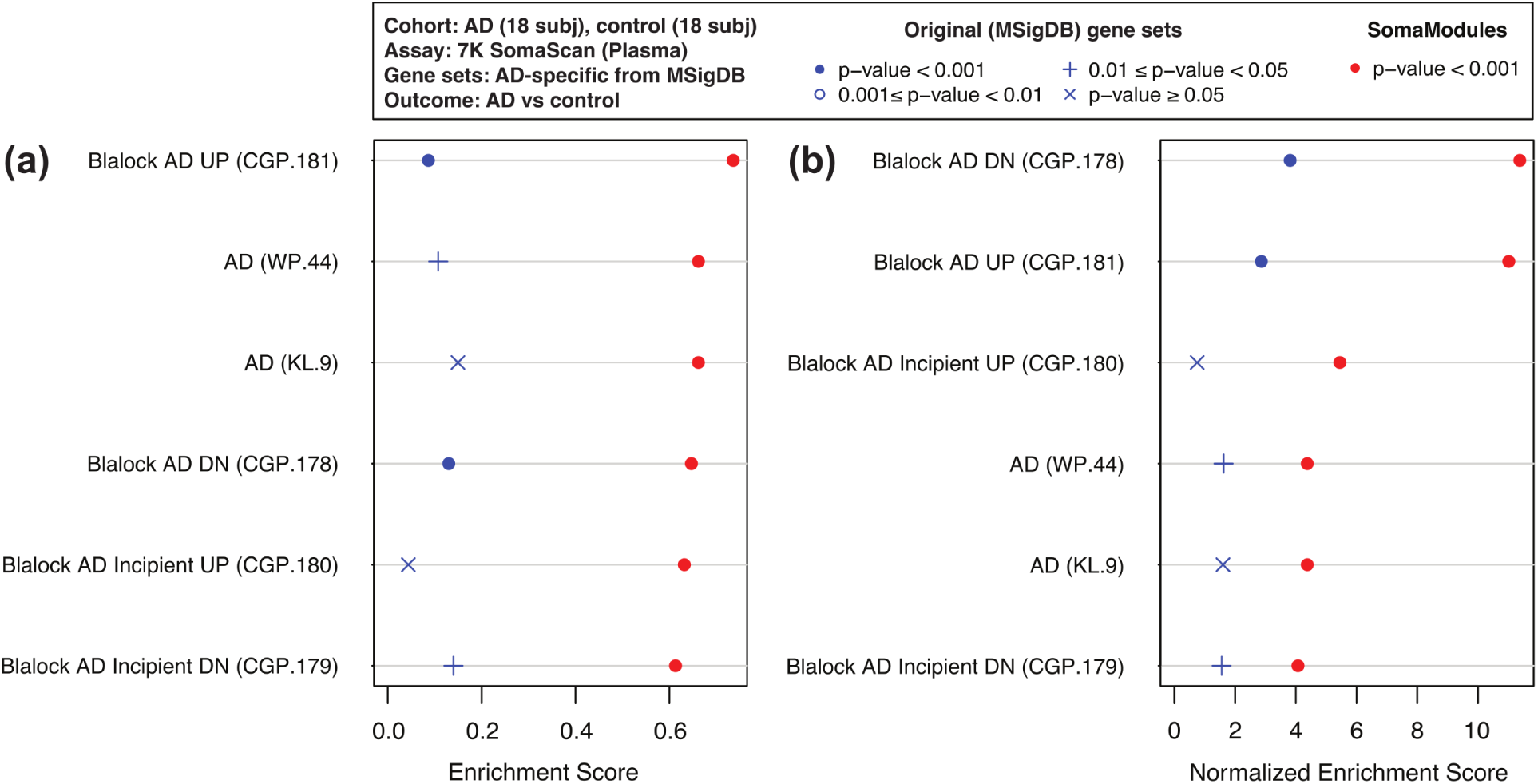
GSEA enrichment scores of AD-specific pathways using 7K SomaScan data from AD vs control plasma samples. Two metrics of pathway enrichment are shown: **(a)** Enrichment Scores; and **(b)** Normalized Enrichment Scores (designed to account for differences in gene set size). For each pathway, SomaModules (red symbols) are compared to the original MSigDB counterparts (blue symbols). Pathways are ordered, from top to bottom, in decreasing order of SomaModule enrichment. Pathways shown are enriched in controls relative to AD patients. Pathway names from MSigDB are followed, in parenthesis, by the identifiers used in our repository, which are formed by a collection prefix (“WP” for WikiPathways, “KL” for KEGG Legacy, and “CGP” for Chemical and Genetic Perturbations) and an integer suffix.

**Figure 4:**
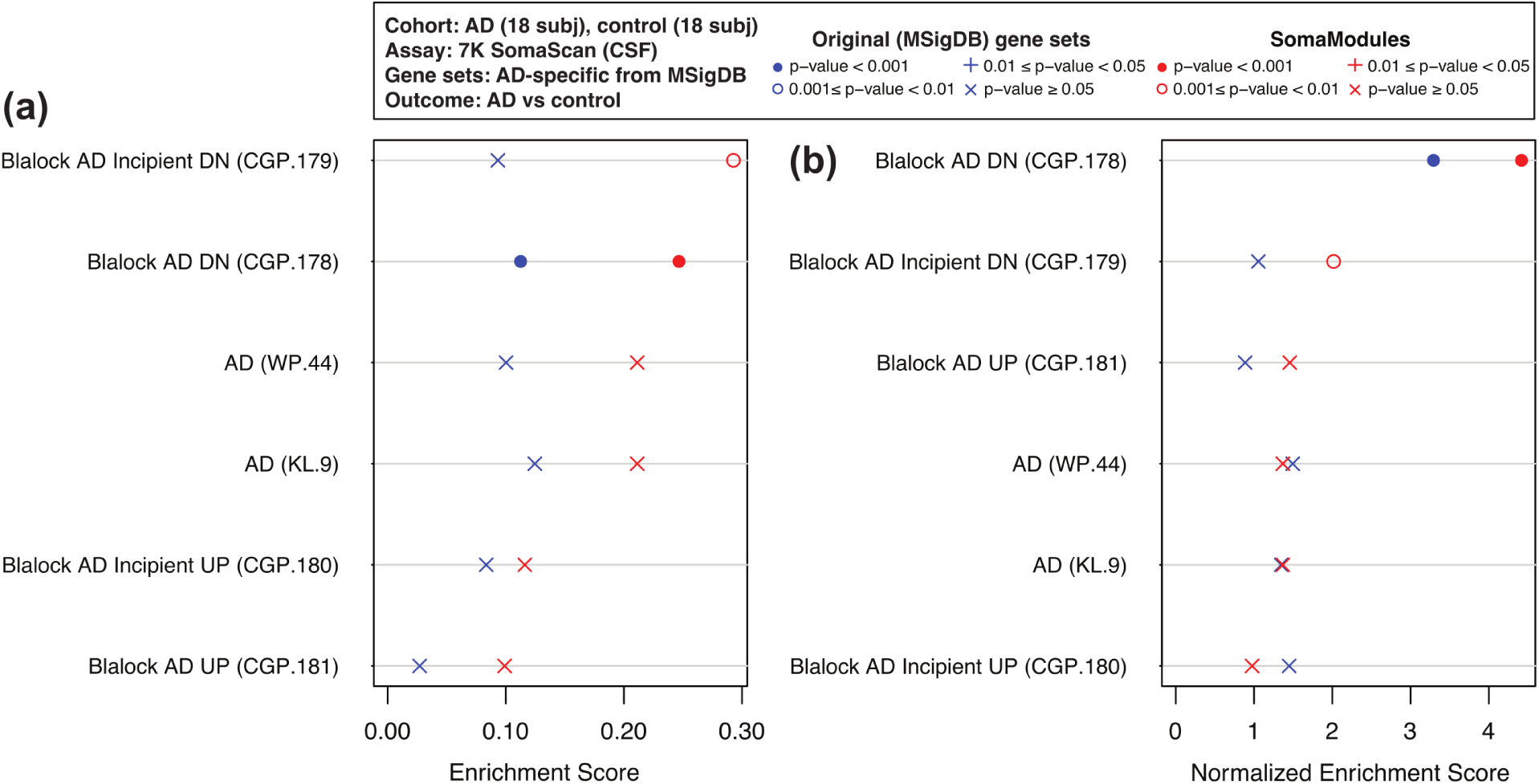
GSEA enrichment scores of AD-specific pathways using 7K SomaScan data from AD vs control CSF samples. Two metrics of pathway enrichment are shown: **(a)** Enrichment Scores; and **(b)** Normalized Enrichment Scores (designed to account for differences in gene set size). For each pathway, SomaModules (red symbols) are compared to the original MSigDB counterparts (blue symbols). Pathways are ordered, from top to bottom, in decreasing order of SomaModule enrichment. Pathways shown are enriched in AD patients relative to controls. Pathway names from MSigDB are followed, in parenthesis, by the identifiers used in our repository, which are formed by a collection prefix (“WP” for WikiPathways, “KL” for KEGG Legacy, and “CGP” for Chemical and Genetic Perturbations) and an integer suffix.

The second example is a longitudinal analysis of physical performance in the BLSA study. Notice that, while SomaModules were built using only the first available visit for each participant (666 samples), for this analysis we used all available visits from those same participants (2542 samples). Using mixed effects models to control for repeat visits, age and sex, we generated rank-ordered lists of 11K SOMAmers associated with 15 different metrics of physical performance. Based on previous findings on the relation between exercise and mitochondrial function in aging, ^71–73^ we hypothesized that mitochondrial pathway enrichment would be significantly associated with physical performance outcomes.

Derived from MitoCarta,^64^ we identified 20 mitochondrial pathways that had one SomaModule counterpart; utilizing those 40 gene sets, we ran separate GSEA analyses for each physical performance outcome. Fig. 5 shows a paired comparison of Enrichment Scores (panel (a)) and Normalized Enrichment Scores (panel (b)) using 6 meter walking time as outcome. We find that, for all the mitochondrial pathways considered, SomaModules (red symbols) are more enriched than their original gene set counterparts from MitoCarta (blue symbols). In fact, while all SomaModules are very significantly enriched (p-value *<* 10^−3^), we observe that the enrichment significance for many of the original gene sets is weaker (10^−3^ ≤ p-value *<* 0.05) or non-significant altogether (p-value ≥ 0.05). Results similarly obtained by considering other physical performance outcomes are provided in Supplementary Data 3 for all choices of GSEA enrichment statistics. In order to compare the absolute values of enrichment between original MitoCarta gene sets and SomaModules derived from them, we performed paired Wilcoxon and Student’s t-tests for each physical performance outcome. Fig. 6 shows the −*log*_10_ (p-value) obtained from paired Wilcoxon tests as a function of the median of paired differences in the absolute value of enrichment scores. In all cases, absolute enrichment scores of SomaModules were greater than the corresponding ones for the MitoCarta pathways of origin; moreover, we observe that, except for normalized enrichment scores using the “narrow walk ratio” and “expanded physical performance battery score” outcomes, all remaining comparisons yielded statistically significant differences. Supplementary Fig. 4 shows similar results using Student’s t-test for comparison. Altogether, these results show that SomaModules have significantly greater enrichment than the original gene set counterparts. These findings are robust and not significantly affected by the choices of enrichment metric or the Kolmogorov enrichment statistic used in the GSEA procedure.

**Figure 5:**
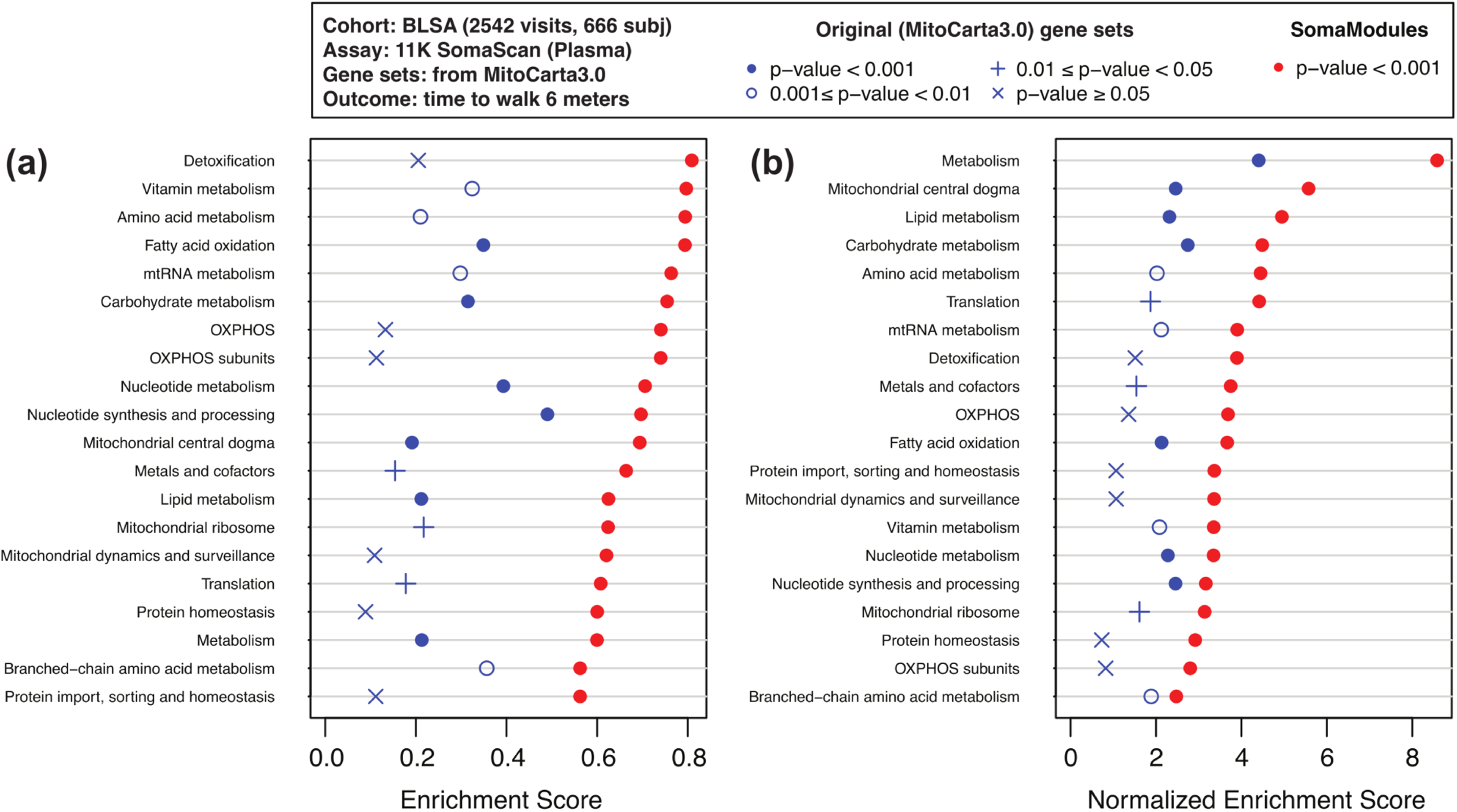
GSEA enrichment scores of mitochondrial pathways associated with the time to walk 6 meters using 11K SomaScan data from 2542 BLSA plasma samples. For each pathway, SomaModules (red symbols) are compared to the original MSigDBS counterparts (blue symbols). Pathways are ordered, from top to bottom, in decreasing order of SomaModule enrichment. **(a)** Enrichment Scores. **(b)** Normalized Enrichment Scores (designed to account for differences in gene set size).

**Figure 6:**
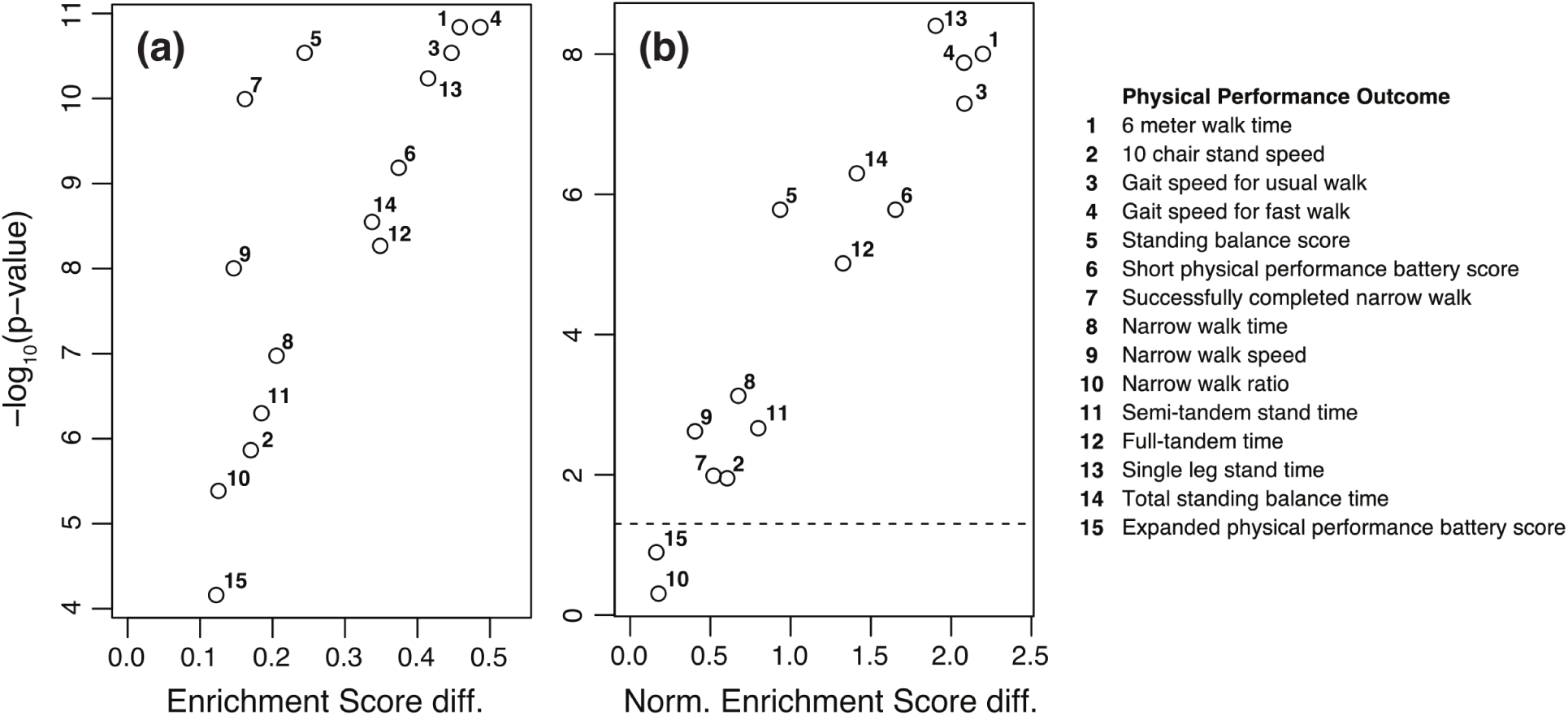
Paired Wilcoxon test significance of enrichment score differences between SomaModules and original MitoCarta pathways for different physical performance outcomes. Panels show differences in **(a)** Enrichment Scores and **(b)** Normalized Enrichment Scores. Physical performance outcomes are listed on the right-hand side. The horizontal dashed line corresponds to p-value = 0.05.

## Conclusions

Given the lack of adequate tools to perform pathway enrichment analysis on proteomic data, this work presents an approach specifically tailored to the SomaScan assay. Using existing, annotated gene sets as the starting point, our framework implements a top-down procedure to identify strongly correlated SOMAmer modules, termed “SomaModules”, based on 11K SomaScan data recently generated from plasma samples in the BLSA study.

The SomaModule framework is a greedy approach to identify, through successive iterations, the largest non-overlapping modules whose median correlation of intra-module SOMAmer pairs is above a predefined threshold value. Throughout this paper, we utilize SomaModules that were generated by using a minimum size of 10 SOMAmers and a minimum intra-module median correlation of 0.5. However, we provide well-documented code to allow users to build their own SomaModule repositories using other threshold criteria of their choice, as well as other pathway sources (including their own, customized gene sets). In this work, we generated two repositories: the first one, based on the latest MSigDB release, contains a total of 41,705 SOMAmer-based gene sets ranging in size from 10 to 1697 SOMAmers; the second one, based on the latest MitoCarta release, contains 80 mitochondrial gene sets ranging in size from 10 to 317 SOMAmers. These repositories can be utilized in any non-topology-based pathway enrichment analysis tool, such as Over-Representation Analysis and Functional Class Scoring (FCS) methods. In this work, we demonstrated the use of our repositories with Gene Set Enrichment Analysis (GSEA), the pioneering and arguably most popular FCS tool. Following the same formatting used in MSigDB, our repositories can be seamlessly integrated to GSEA.

We validated our repositories with two case examples. In the first one, we used 7K SomaScan data from a study of Alzheimer’s Disease (AD) patients and non-AD controls to test AD-specific pathways from MSigDB. We found that SomaModules had significantly higher enrichment than the original gene set counterparts. In our second case example, we used 11K SomaScan data from the BLSA to explore the enrichment of mitochondrial pathways from MitoCarta in association with a variety of physical performance outcomes. In agreement with our previous example, we found that SomaModules had greater enrichment than the original gene set counterparts and that those differences were generally significant. These findings were robust, not significantly affected by the choice of enrichment metric or the Kolmogorov enrichment statistic used in the GSEA procedure. Comparisons performed with paired Wilcoxon and Student’s t-tests consistently showed that SomaModules had significantly greater enrichment than the original gene set counterparts.

As SomaScan becomes more widely adopted as a state-of-the-art tool for proteomics discovery, we hope that this work will serve as a valuable technical reference and resource for the growing user community. We finally note that similar procedures may be adopted for follow-up work extending annotated pathways (or uncovering novel ones) based on data from other proteomics platforms, such as Olink and targeted mass spectrometry, and even other omics assays beyond proteomics.^30^

## Acknowledgement

This research was supported entirely by the Intramural Research Program of the National Institute on Aging, NIH. The authors thank Cassandra Blew and Cassandra Joynes for useful discussions and preliminary analyses of SomaModule characteristics.

## Supporting Information Available

The following files are available online:

- Supplementary Table 1: Number of gene sets in each MSigDB collection, derived using different threshold combinations for minimum gene set size and intra-cluster correlation.
- Supplementary Table 2: Summary of results from WGCNA runs using different sets of parameters.
- Supplementary Figure 1: Mean correlation density distributions for parent-child SomaModule pairs derived from different MSigDB collections.
- Supplementary Figure 2: WGCNA grid-search for the optimal soft-thresholding power (beta) for network construction.
- Supplementary Figure 3: Volcano plots showing differentially abundant SOMAmers from an Alzheimer’s Disease vs control study using 7K SomaScan.
- Supplementary Figure 4: Paired Student’s t-test significance of enrichment score differences between SomaModules and original MitoCarta pathways for different physical performance outcomes.
- Supplementary Data 1: GSEA enrichment scores of AD-specific pathways using 7K SomaScan data from AD vs control plasma samples.
- Supplementary Data 2: GSEA enrichment scores of AD-specific pathways using 7K SomaScan data from AD vs control CSF samples.
- Supplementary Data 3: GSEA enrichment scores of mitochondrial pathways associated with 15 physical performance metrics using 11K SomaScan data from 2542 BLSA plasma samples.

## For Table of Contents Only

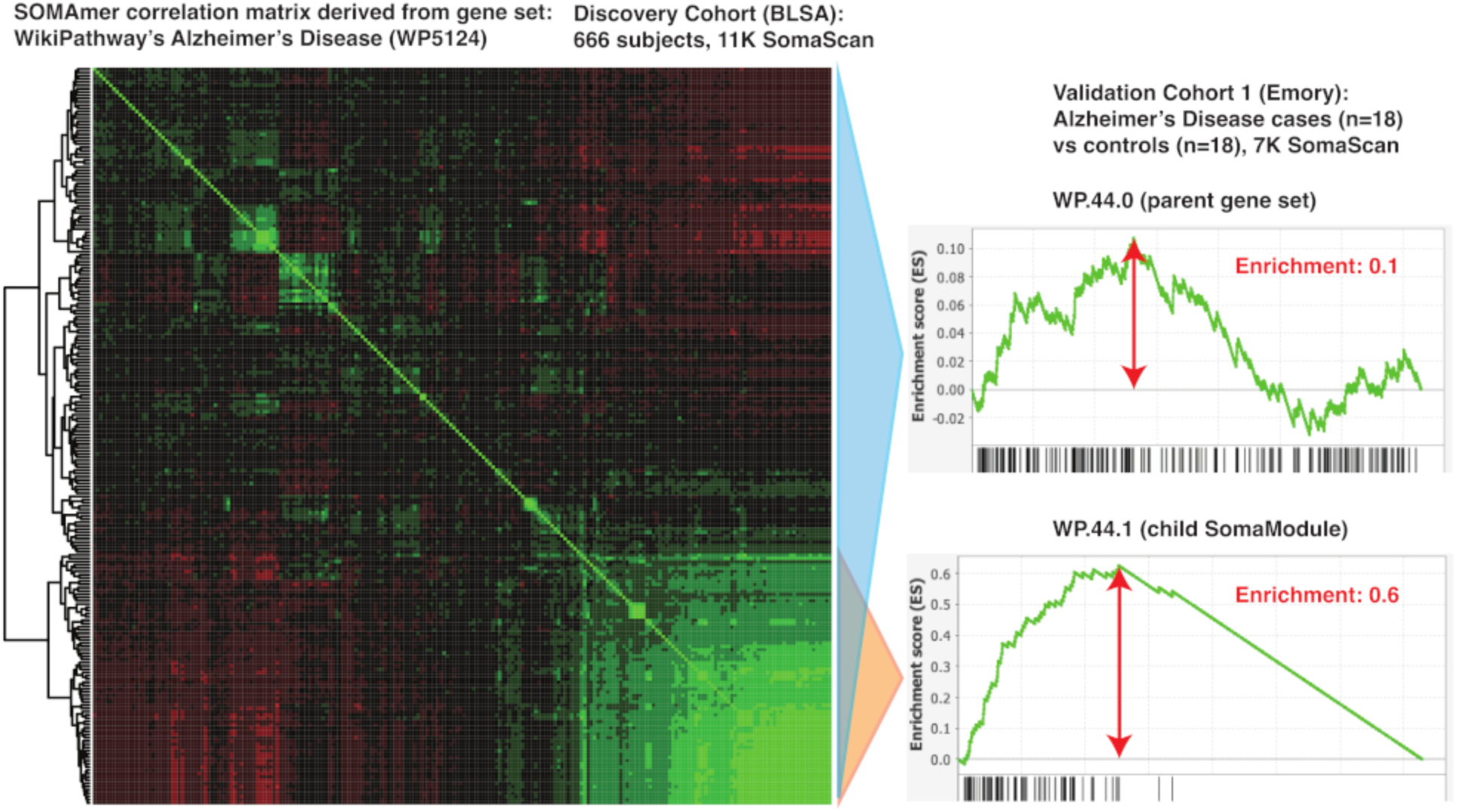

