## Supplementary Tables for "SomaModules: a pathway enrichment approach tailored to SomaScan data"

| ID | Description | MSigDB ID | n = 5 | | | n = 10 | | | n = 20 | | |
|---|---|---|---|---|---|---|---|---|---|---|---|
| | | | r = 0.3 | r = 0.5 | r = 0.7 | r = 0.3 | r = 0.5 | r = 0.7 | r = 0.3 | r = 0.5 | r = 0.7 |
| H | Hallmarks | h.all | 152 | 94 | 58 | 123 | 86 | 58 | 94 | 79 | 55 |
| POS | Positional | c1.all | 499 | 373 | 268 | 349 | 304 | 253 | 268 | 261 | 243 |
| CGP | Chemical and Genetic Perturbations | c2.cgp | 7480 | 5066 | 3561 | 5581 | 4318 | 3346 | 4305 | 3797 | 3231 |
| BC | BioCarta | c2.cp.biocarta | 405 | 353 | 283 | 293 | 281 | 254 | 234 | 232 | 224 |
| K | KEGG | c2.cp.kegg_medicus | 621 | 562 | 437 | 431 | 395 | 367 | 366 | 362 | 361 |
| KL | KEGG Legacy | c2.cp.kegg_legacy | 423 | 318 | 222 | 305 | 272 | 210 | 247 | 234 | 199 |
| PID | Pathway Interaction Database | c2.cp.pid | 441 | 365 | 289 | 344 | 323 | 273 | 260 | 252 | 228 |
| R | Reactome | c2.cp.reactome | 2858 | 2226 | 1657 | 2106 | 1827 | 1507 | 1696 | 1562 | 1381 |
| WP | WikiPathways | c2.cp.wikipathways | 1531 | 1152 | 889 | 1108 | 973 | 819 | 875 | 824 | 757 |
| MIR | microRNA Targets | c3.mir.mirdb | 6261 | 4234 | 2757 | 4719 | 3666 | 2605 | 3468 | 3009 | 2423 |
| MIRL | microRNA Targets Legacy | c3.mir.mir_legacy | 580 | 407 | 275 | 464 | 365 | 262 | 341 | 304 | 238 |
| TFT | Transcription Factor Targets | c3.tft.gtrd | 1273 | 858 | 510 | 1092 | 789 | 493 | 931 | 730 | 475 |
| TFTL | Transcription Factor Targets Legacy | c3.tft.tft_legacy | 1952 | 1218 | 761 | 1602 | 1146 | 743 | 1192 | 1031 | 716 |
| 3CA | Curated Cancer Cell Atlas | c4.3ca | 361 | 243 | 174 | 223 | 184 | 157 | 162 | 154 | 151 |
| CGN | Cancer Gene Neighborhoods | c4.cgn | 1038 | 776 | 552 | 798 | 670 | 522 | 599 | 543 | 471 |
| CM | Cancer Modules | c4.cm | 973 | 647 | 491 | 747 | 561 | 455 | 579 | 478 | 424 |
| GOBP | GO Biological Process | c5.go.bp | 11224 | 8200 | 6103 | 8351 | 6970 | 5672 | 6880 | 6214 | 5402 |
| GOCC | GO Cellular Component | c5.go.cc | 1312 | 954 | 688 | 1020 | 823 | 649 | 833 | 729 | 615 |
| GOMF | GO Molecular Function | c5.go.mf | 1877 | 1431 | 1104 | 1414 | 1215 | 1019 | 1176 | 1078 | 968 |
| HPO | Human Phenotype Ontology | c5.hpo | 8350 | 6218 | 4391 | 6441 | 5337 | 4150 | 5178 | 4679 | 3979 |
| ONC | Oncogenic | c6.all | 579 | 336 | 213 | 437 | 295 | 204 | 287 | 233 | 194 |
| IMM | Immunologic | c7.immunesigdb | 15211 | 9931 | 5967 | 12179 | 9363 | 5931 | 9381 | 8583 | 5695 |
| VAX | Vaccine Response | c7.vax | 573 | 406 | 304 | 458 | 365 | 296 | 373 | 343 | 290 |
| CT | Cell Type | c8.all | 2208 | 1305 | 898 | 1694 | 1177 | 878 | 1296 | 1070 | 853 |
| **TOTAL** | | | 68182 | 47673 | 32852 | 52279 | 41705 | 31123 | 41021 | 36781 | 29573 |

**Supplementary Table 1.** Number of gene sets in each MSigDB collection, derived using different threshold combinations for minimum gene set size (n = 5, 10, 20) and minimum intra-cluster correlation (r = 0.3, 0.5, 0.7). Results for the threshold values used in this paper (n=10, r=0.5) are highlighted in grey, which match those reported in Table 1.

| power | deepSplit | TOMType | networkType | n_mod | n_soma_mod | n_soma | PC1_var_expl |
|---|---|---|---|---|---|---|---|
| 5 | 1 | none | signed | 2 | 94\|44 | 138 | 0.329 |
| 5 | 1 | none | signed hybrid | 2 | 98\|40 | 138 | 0.351 |
| 5 | 1 | unsigned | signed | 2 | 94\|44 | 138 | 0.329 |
| 5 | 1 | unsigned | signed hybrid | 3 | 84\|44\|10 | 138 | 0.416 |
| 5 | 1 | signed | signed | 2 | 94\|44 | 138 | 0.329 |
| 5 | 1 | signed | signed hybrid | 3 | 84\|44\|10 | 138 | 0.416 |
| 5 | 2 | none | signed | 2 | 94\|44 | 138 | 0.329 |
| 5 | 2 | none | signed hybrid | 3 | 88\|40\|10 | 138 | 0.43 |
| 5 | 2 | unsigned | signed | 3 | 81\|44\|13 | 138 | 0.369 |
| 5 | 2 | unsigned | signed hybrid | 3 | 84\|44\|10 | 138 | 0.416 |
| 5 | 2 | signed | signed | 3 | 81\|44\|13 | 138 | 0.369 |
| 5 | 2 | signed | signed hybrid | 3 | 84\|44\|10 | 138 | 0.416 |
| 5 | 3 | none | signed | 4 | 69\|43\|14\|12 | 138 | 0.363 |
| 5 | 3 | none | signed hybrid | 3 | 88\|40\|10 | 138 | 0.43 |
| 5 | 3 | unsigned | signed | 3 | 81\|44\|13 | 138 | 0.369 |
| 5 | 3 | unsigned | signed hybrid | 3 | 84\|44\|10 | 138 | 0.416 |
| 5 | 3 | signed | signed | 3 | 81\|44\|13 | 138 | 0.369 |
| 5 | 3 | signed | signed hybrid | 3 | 84\|44\|10 | 138 | 0.416 |
| 5 | 4 | none | signed | 6 | 43\|43\|17\|12\|12\|11 | 138 | 0.337 |
| 5 | 4 | none | signed hybrid | 3 | 88\|40\|10 | 138 | 0.43 |
| 5 | 4 | unsigned | signed | 3 | 81\|44\|13 | 138 | 0.369 |
| 5 | 4 | unsigned | signed hybrid | 3 | 84\|44\|10 | 138 | 0.416 |
| 5 | 4 | signed | signed | 3 | 81\|44\|13 | 138 | 0.369 |
| 5 | 4 | signed | signed hybrid | 3 | 84\|44\|10 | 138 | 0.416 |

**Supplementary Table 2.** Summary of results from WGCNA runs using different sets of parameters: n_mod = number of modules; n_soma_mod = number of SOMAmers in each module; n_soma = number of SOMAmers across all modules; PC1_var_expl = variance explained by Principal Component 1 averaged over all modules. Highlighted in grey are the runs that maximize PC1_var_expl (they all correspond to the same solution).
