## Supplementary Figures for "SomaModules: a pathway enrichment approach tailored to SomaScan data"

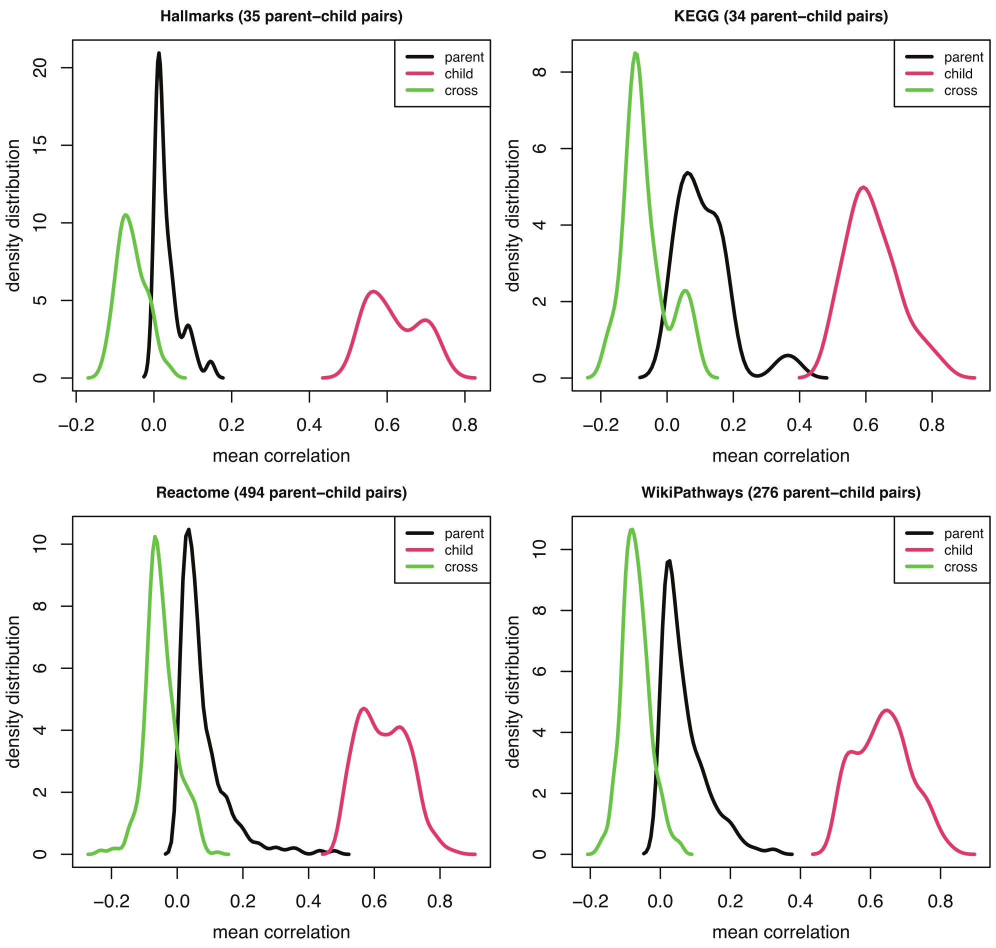


**Supplementary Figure 1: Mean correlation density distributions for parent-child SomaModule pairs derived from different MSigDB collections, as indicated.**

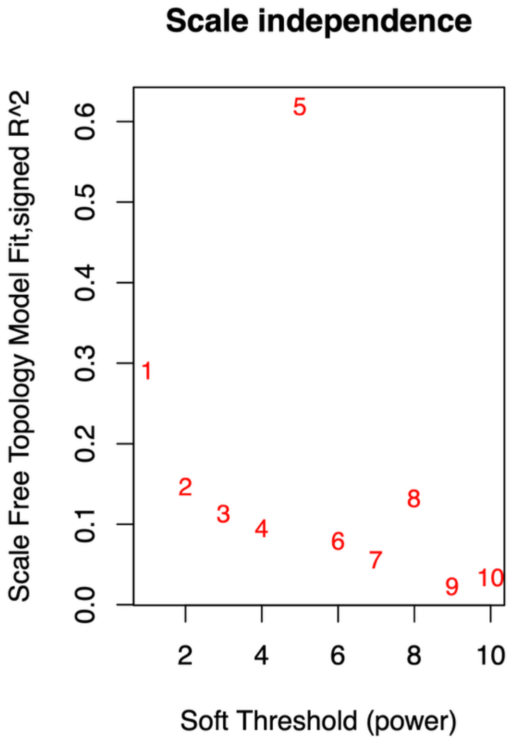


**Supplementary Figure 2: WGCNA grid-search for the optimal soft-thresholding power (beta) for network construction.**

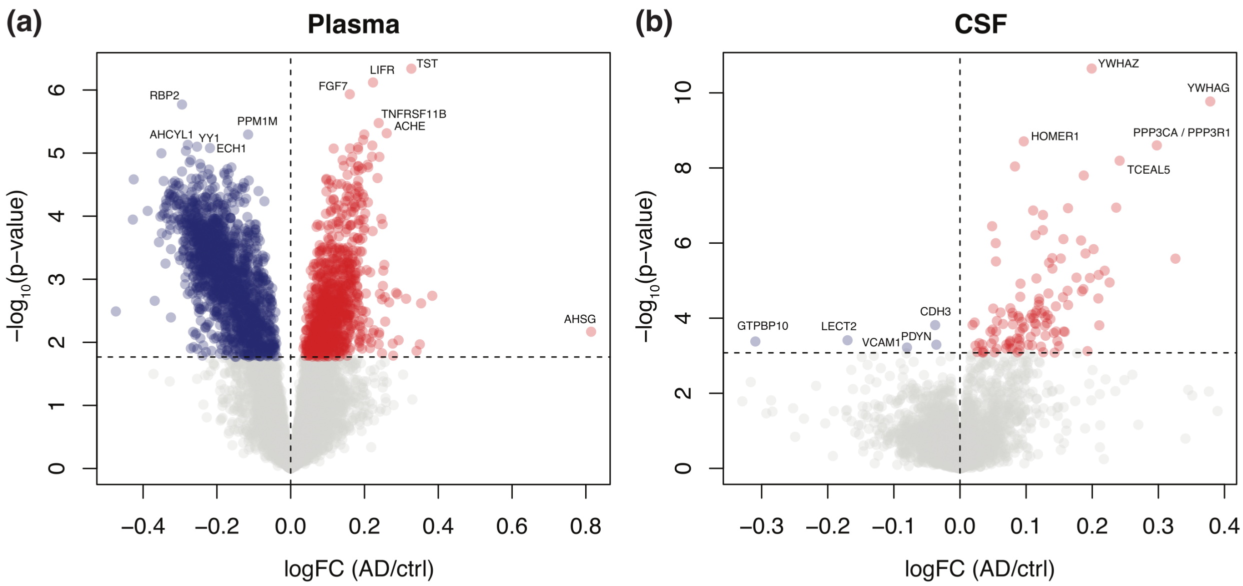


**Supplementary Figure 3: Volcano plots showing differentially abundant SOMAmers from an Alzheimer's Disease (AD) vs control study using 7K SomaScan.** The horizontal dashed lines show the Benjamini-Hochberg adjusted p-value=0.05. Red symbols show SOMAmers significantly over-abundant in AD relative to controls; blue symbols correspond to SOMAmers significantly over-abundant in controls relative to AD patients. **(a)** Plasma samples ($n_{red}$=923, $n_{blue}$=1562). **(b)** Cerebrospinal fluid (CSF) samples ($n_{red}$=117, $n_{blue}$=5).

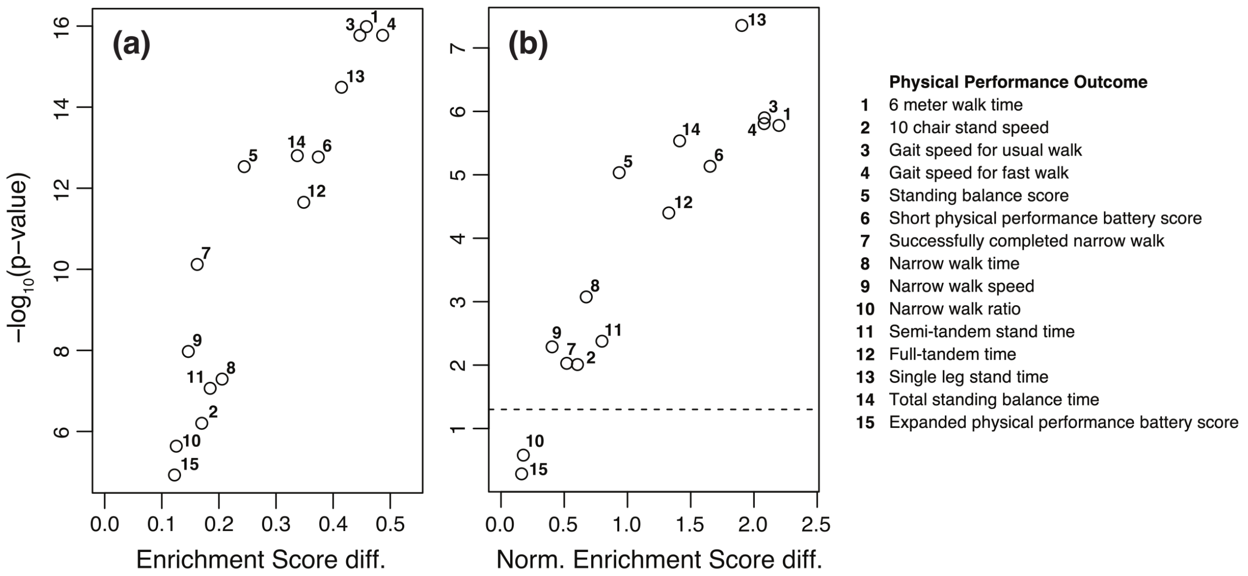


**Supplementary Figure 4: Paired Student's t-test significance of enrichment score differences between SomaModules and original MitoCarta pathways for different physical performance outcomes.** Panels show differences in **(a)** Enrichment Scores and **(b)** Normalized Enrichment Scores. Physical performance outcomes are listed on the right-hand side. The horizontal dashed line corresponds to p−value = 0.05.
